# Resistance to serine in *Bacillus subtilis*: Identification of the serine transporter YbeC and of a metabolic network that links serine and threonine metabolism

**DOI:** 10.1101/2020.05.20.106443

**Authors:** Anika Klewing, Byoung Mo Koo, Larissa Krüger, Anja Poehlein, Daniel Reuß, Rolf Daniel, Carol A. Gross, Jörg Stülke

## Abstract

The Gram-positive bacterium *Bacillus subtilis* uses serine not only as building block for proteins but also as an important precursor in many anabolic reactions. Moreover, a lack of serine results in the initiation of biofilm formation. However, in excess serine inhibits the growth of *B. subtilis*. To unravel the underlying mechanisms, we isolated suppressor mutants that can tolerate toxic serine concentrations by three targeted and non-targeted genome-wide screens. All screens as well as genetic complementation in *Escherichia coli* identified the so far uncharacterized permease YbeC as the major serine transporter of *B. subtilis*. In addition to YbeC, the threonine transporters BcaP and YbxG make minor contributions to serine uptake. A strain lacking these three transporters was able to tolerate 100 mM serine whereas the wild type strain was already inhibited by 1 mM of the amino acid. The screen for serine-resistant mutants also identified mutations that result in increased serine degradation and in increased expression of threonine biosynthetic enzymes suggesting that serine toxicity results from interference with threonine biosynthesis.

**Originality-Significance Statement:** Serine is an important precursor for many biosynthetic reactions, and lack of this amino acid can induce biofilm formation in *Bacillus subtilis*. However, serine is toxic for the growth of *B. subtilis*. To understand the reason(s) for this toxicity and to identify the so far unknown serine transporter(s) of this bacterium, we performed exhaustive mutant screens to isolate serine-resistant mutants. This screen identified YbeC, the major serine transporter of *B. subtilis*. Moreover, we observed an intimate link between serine and threonine metabolism that is responsible for serine toxicity by inhibiting threonine biosynthesis.

## Introduction

As building block of proteins, amino acids are central to the physiology of any living cell. In addition to their role as substrates in protein biosynthesis, they can be used as carbon and nitrogen sources. Moreover, some amino acids are also required for bacterial cell wall biosynthesis and as osmoprotectants. Accordingly, the acquisition of amino acids is an essential task of all cells. This can be achieved by the direct uptake of amino acids present in the growth medium, by the uptake and intracellular degradation of peptides and by *de novo* biosynthesis. Many bacteria such as the model organisms *Escherichia coli* and *Bacillus subtilis* are capable of synthesizing all amino acids whereas others such as the minimal bacteria of the genus *Mycoplasma* completely depend on the uptake of amino acids.

While amino acids are essential for the cells, increased concentrations of some amino acids such as glutamate, threonine or serine can be harmful (Lamb and Bott 1979a; Lamb and Bott 1979b; Lachowicz *et al*., 1996; Ogawa *et al*., 1998; Mundhada *et al*., 2017; Belitsky, 2015; Commichau *et al*., 2008; Belitsky and Sonenshein, 1998). Therefore, the homeostasis of the amino acids must be tightly controlled to adjust the intracellular levels of each amino acid to the actual need of the cell. This requires balanced activities of systems for amino acid acquisition and degradation. For the Gram-positive model bacterium *B. subtilis*, amino acid metabolism is one of the few functions in core metabolism that have not yet been completely elucidated. This is the case both for the biosynthetic pathways and for amino acid transport.

The genome of *B. subtilis* encodes 47 known and predicted amino acid transporters (Zhu and Stülke, 2018). For 19 of these transporters, substrates have been identified, and for four additional transporters, tentative substrates can be assigned based on mutant properties (for YbxG; Commichau *et al*., 2015) and on the assignment to particular regulons (AlsT, YvbW, and YvsH; Randazzo *et al*., 2017; Wels *et al*., 2008; Rodionov *et al*., 2003). No functional assignment can so far be made for eleven potential transporters. It is important to note that some proteins that are members of typical amino acid transporter families do actually transport other substrates, such as the recently described potassium transporter KimA (Gundlach *et al*., 2017). A complete overview on the known and potential amino acid transporters of *B. subtilis* can be found in Table S1 (see also http://subtiwiki.uni-goettingen.de/v3/category/view/SW%201.2, Zhu and Stülke, 2018). Importantly, no transporters have so far been identified or proposed for alanine, glycine, serine, asparagine, and the aromatic amino acids phenylalanine and tyrosine. The identification of new amino acid transporters is hampered by two peculiarities: For one amino acid, there are often multiple transporters, as has been shown for arginine, proline, or the branched-chain amino acids (Calogero *et al*., 1994; Gardan *et al*., 1995; Sekowska *et al*., 2001; Zaprasis *et al*., 2014; Belitsky, 2015). On the other hand, many permeases have a relatively weak substrate specificity, i. e. they are able to transport multiple amino acids, as shown for BcaP or GltT (Belitsky 2015; Zaprasis *et al*., 2015).

We are interested in the identification of the functions that are required to sustain the life of a minimal cell and in the corresponding set of genes and proteins. In an analysis of the genome of *B. subtilis*, amino acid transporters were proposed to be kept in a minimal genome rather than biosynthetic genes, since this would require fewer genes (Reuss *et al*., 2016). However, as indicated above, no transporters have been identified for several amino acids. Accordingly, biosynthetic pathways were included for those amino acids. A minimal organism capable of transporting amino acids but not to produce them is expected to be viable on complex media but would be unable to grow on minimal medium in the absence of added amino acids. As minimal bacterial strains have a huge potential for biotechnological applications (Suárez *et al*., 2019), the ability to produce amino acids may be important for growth on cheap minimal salts substrates.

Serine is an important amino acid because this molecule is a not only a building block for protein synthesis but also a precursor of nucleotides, phospholipids, redox molecules, and other amino acids. In addition, decreased level of intracellular serine can be a signal for initiation of biofilm formation in *B. subtilis* (Subramanian *et al*., 2013), suggesting that regulation of serine homeostasis is very important. However, the metabolism of serine is not yet completely understood in *B. subtilis*. For this amino acid, no transporter has been identified, and the knowledge of biosynthetic pathways has remained limited until recently. Indeed, the serine biosynthesis pathway has been completed just recently in the frame of a genome-scale deletion study by the identification of the SerB phosphoserine phosphatase that catalyzes the last step of the pathway (Koo *et al*., 2017) (see Fig. 1 for an overview on serine metabolism in *B. subtilis*). Moreover, the reasons for serine toxicity have remained enigmatic. In *E. coli*, it has been suggested that serine binds and inactivates the bifunctional enzyme aspartate kinase/homoserine dehydrogenase (ThrA) (Mundhada *et al*., 2017) thus interfering with threonine biosynthesis.

**Fig. 1.**
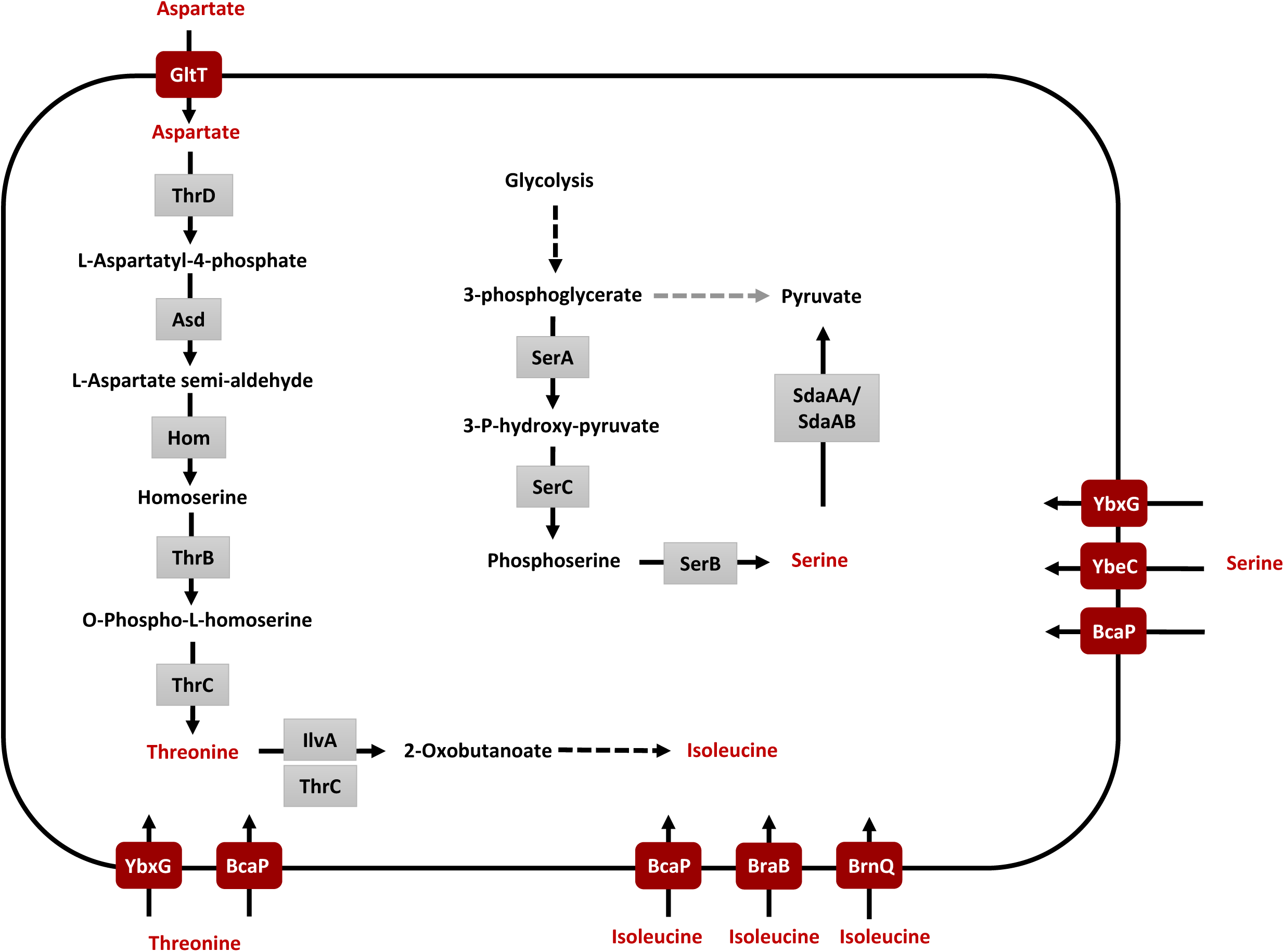
Serine and threonine metabolic pathways in *B. subtilis*. The model shows the relevant transporters, the biosynthesis of threonine, and its role as precursor for isoleucine biosynthesis as well as the pathways for serine biosynthesis and degradation.

In this work, we have taken advantage of the serine toxicity phenotype of *B. subtilis* to devise reverse and forward genetic screens to identify serine transporters. Our analysis identified YbeC as the major serine transporter, and clarified the roles of threonine transporters in uptake of serine. Using a suppressor screen, we also isolated metabolic mutants that circumvent serine toxicity. These mutants exhibit either more efficient serine degradation or overexpression of genes for the threonine and isoleucine biosynthetic pathways suggesting that one or more enzymes in this pathway are inhibited by serine.

## Results

### Overview of the genetic approaches used in this work

All of our screens and selections were based on the fact that addition of serine to minimal medium is toxic for *B. subtilis* 168, whereas the addition of serine to complex LB medium did not interfere with growth of the bacteria. Moreover, the addition of specific amino acids such as threonine to minimal medium also overcomes serine toxicity (Vandeyar and Zahler, 1986; Lachowicz *et al*., 1996). These observations suggest that intracellular serine interferes with amino acid metabolism. We might therefore expect that strains would become resistant to serine toxicity either by eliminating the major serine transporter, or by altering amino acid metabolism of genes related to serine toxicity. To identify these genes, we used the following approaches:

#### 1. A targeted screen for transporters

For this purpose, we chose twelve candidate transporters that met two criteria: First, these transporters have been poorly studied in *B. subtilis*, and second, they are expressed during vegetative growth. These transporters are AapA, AlsT, MtrA, SteT, YbeC, YbgF, YdgF, YecA, YodF, and YtnA. Mutants for the corresponding genes (see Table S2) were constructed and analysed for the ability to grow in the presence of 244 µM L-serine. While all strains were capable of growing on minimal medium in the absence of serine, only the *ybeC* mutant strain GP1886 was able to grow in the presence of serine, suggesting that YbeC might act as serine transporter.

#### 2. A suppressor screen aimed at identifying mutants altered in related amino acid metabolism

We selected for loss of serine toxicity in the wild type strain and in a Δ*serA* mutant that is auxotrophic for serine strain and depends on the uptake of serine for growth. Of eight studied suppressor strains, four were transporter (*ybeC*) mutants and the remaining strains exhibited genetic lesions related to amino acid metabolism. Interestingly, the *ybeC* mutation was also found in the *serA* mutant as a suppressor, indicating that serine can be transported in *ybeC* mutant.

#### 3. A screen of the entire *B. subtilis* deletion library for loss of serine toxicity

In order to make sure that the screens described above were exhaustive, we also made use of the deletion library that encompasses all non-essential genes of *B. subtilis* (Koo *et al*., 2017). In this library, each reading frame is replaced by an antibiotic cassette with a relatively strong outwardly facing promoter, so resistance to serine toxicity could result from gene deletion or from overexpression of downstream genes. To distinguish between these possibilities, we removed the antibiotic cassette and retested the phenotype. If the phenotype was retained following removal of the antibiotic cassette, then the phenotype was caused by gene deletion; if not it was likely due to overexpression of downstream genes. This screen identified both the transporter, YbeC, and genetic lesions related amino acid metabolism (Table 1).

**Table 1.**
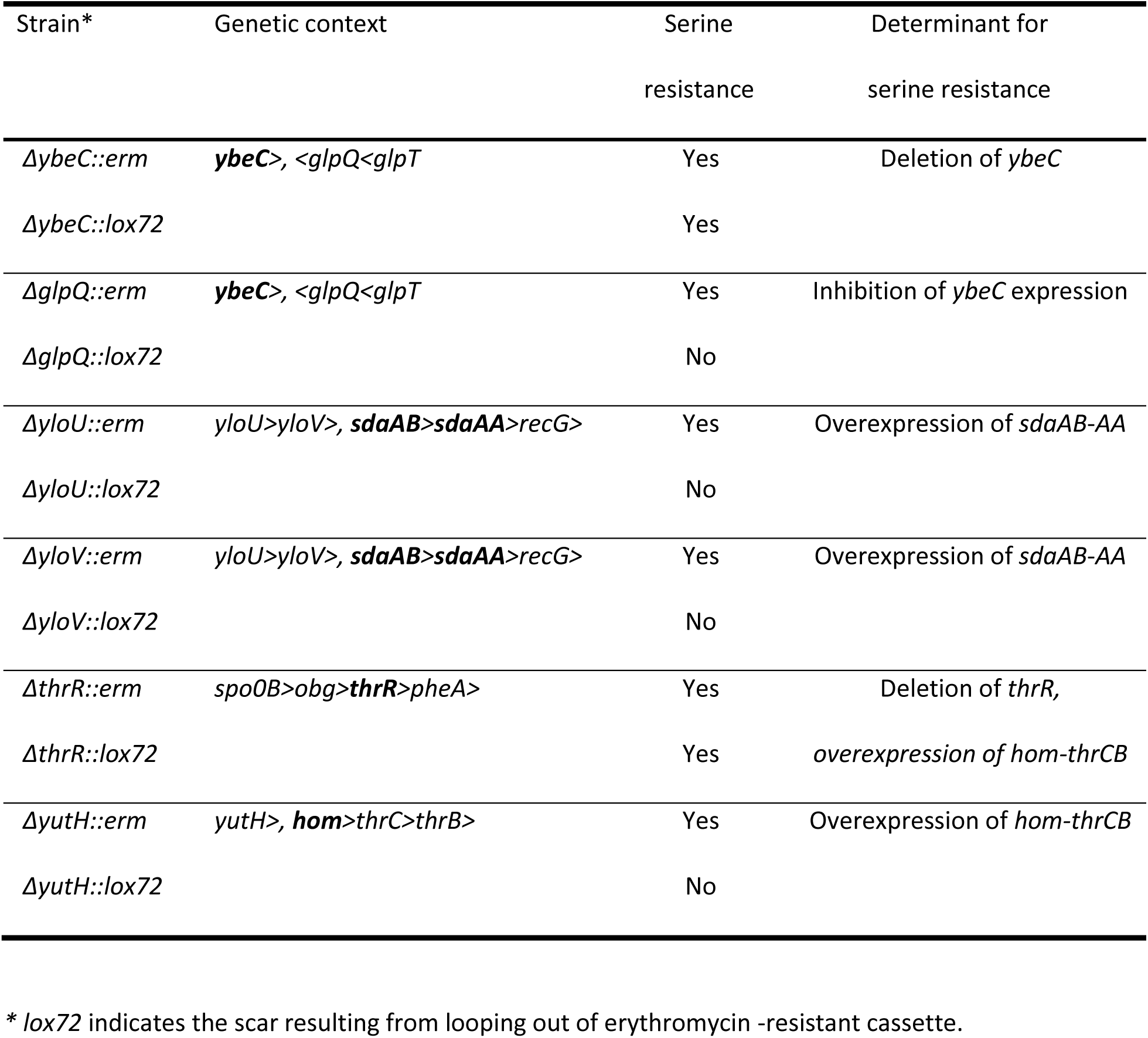
Serine resistant mutants identified from the genome-wide screen

### Identification of a permease that confers sensitivity to L-serine

Both our targeted screen of the 13 expressed, uncharacterized transporters and our screen of the *B. subtilis* deletion library identified only a single putative transporter, YbeC, the loss of which conferred resistance to serine. Supporting the importance of YbeC, 50% of the mutants from the suppressor screen (Selection #2) had mutations targeting *ybeC*. Additionally, the *glpQ* mutant (BKE02130), a strain from the whole genome screen (Selection #3) that suppressed serine toxicity due to overexpression, likely generates an abundant *ybeC* antisense RNA (Table 1). The net effect of antisense expression is to decrease *ybeC* expression, explaining its serine resistance phenotype.

To test whether the *ybeC* mutant is also resistant to higher concentrations of serine, we cultivated the mutant GP1886 at increasing serine concentrations (up to 100 mM), and recorded growth of the bacteria. While the wild type strain was unable to grow at concentrations exceeding 244 µM, the *ybeC* mutant was able to tolerate as much as 11 mM serine (see Table 2). In addition to serine, the anti-metabolite serine hydroxamate also inhibits growth of *B. subtilis*. To test whether loss of YbeC allows growth in the presence of this serine analogue, we cultivated the wild type strain 168 and the *ybeC* mutant GP1886 in the presence of DL-serine hydroxamate (1 mg/ml). As shown in Fig. 2A, the wild type was sensitive to this molecule whereas the *ybeC* mutant was somewhat more resistant. Thus, loss of YbeC confers resistance to both serine and its toxic analogue serine hydroxamate, suggesting that the protein is a serine transporter.

**Table 2:** Resistance of selected *B. subtilis* mutants towards serine.

| Strain | Relevant genotype | Tolerated serine concentration (mM)<sup>1</sup> |
| --- | --- | --- |
| 168 | Wild type | < 1 |
| GP2786 | $\Delta ybeC$ | 11 |
| BKE09460 | $\Delta bcaP$ | 2 |
| GP2396 | $\Delta ybxG$ | 1.5 |
| GP2949 | $\Delta ybeC \Delta bcaP$ | 40 |
| GP2951 | $\Delta ybeC \Delta ybxG$ | 25 |
| GP2952 | $\Delta bcaP \Delta ybxG$ | 4 |
| GP2950 | $\Delta ybeC \Delta bcaP \Delta ybxG$ | 100 |

<sup>1</sup> The tolerated serine concentrations were determined by cultivating the strains in liquid C Glc minimal medium in the presence of different serine concentrations. Note that the results obtained with plates and liquid medium can differ slightly.

**Fig. 2.**
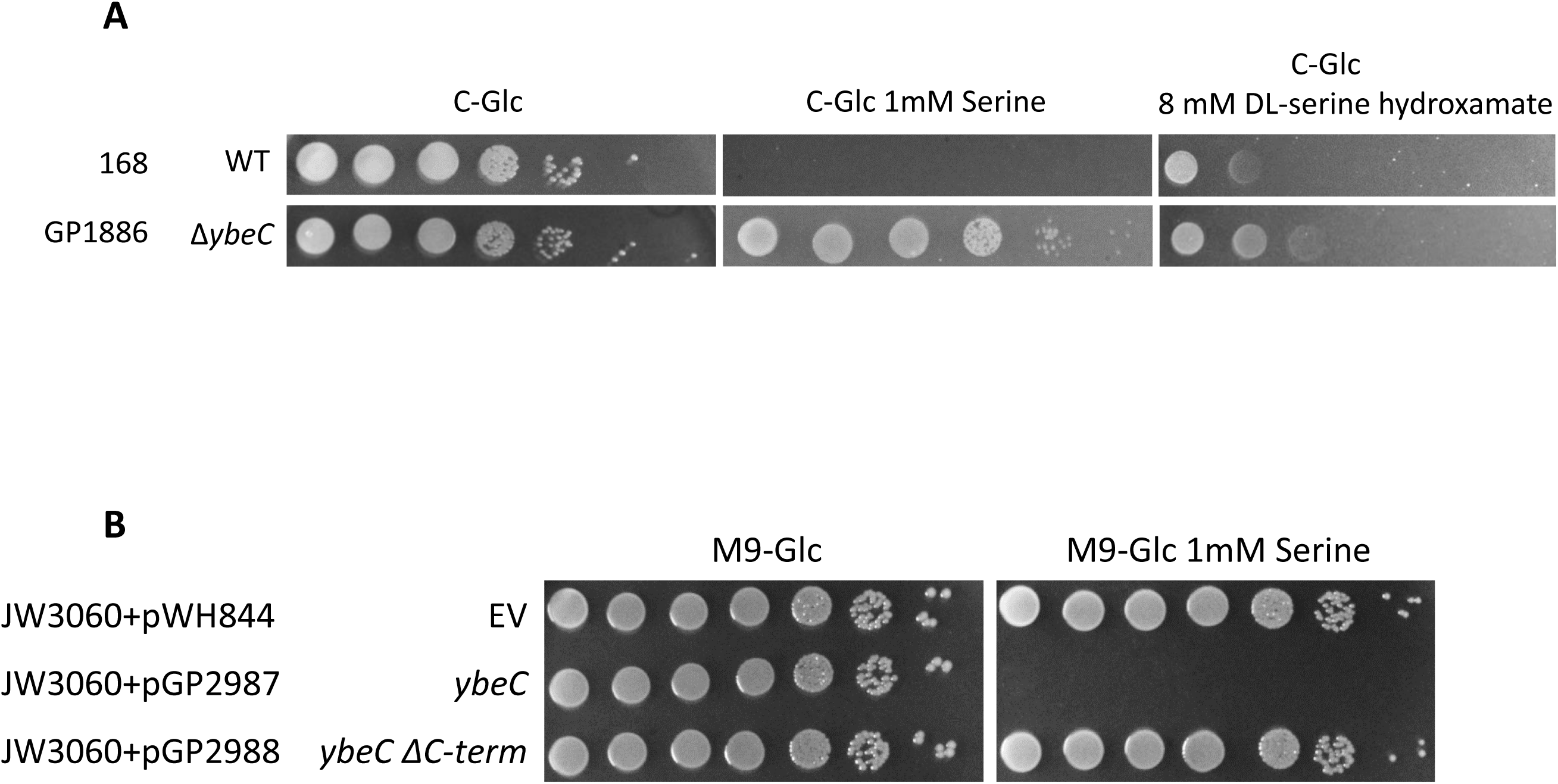
YbeC is a serine transporter. **A.** Sensitivity of the wild type strain 168 and the *ybeC* deletion mutant to serine and the toxic serine analogue DL-serine hydroxamate. Cells of the wild type 168 and the *ybeC* deletion mutant were grown in C-Glc minimal medium to an OD_600_ of 1.0 and serial dilutions (10-fold) were prepared. These samples were plated on C-Glc minimal plates containing no serine, 1 mM serine or 8mM DL-serine hydroxamate. The plates were incubated at 37°C for 48 h. **B.** Serine transport complementation assay in *E. coli*. The growth of the *E. coli sstT* mutant JW3060 harboring the empty vector (pWH844) was compared to the growth of JW3060 with a plasmid encoding the full-length YbeC (pGP2987) or YbeC without the C-terminus (pGP2988) on M9 minimal plates in the presence and absence of serine. The plates were incubated at 37°C for 48 h.

### Isolation and initial characterization of mutants that are able to grow in the presence of serine

The targeted analysis of potential amino acid transporters identified YbeC as the only candidate serine transporter. In an attempt to identify more genes involved in serine toxicity, we cultivated the *B. subtilis* wild type strain 168 in the presence of different concentrations of serine. Moreover, we used the serine auxotrophic *serA* mutant that depends on the uptake of serine for growth, and the *ybeC* mutant that already tolerates up to 11 mM of serine (see above). In total, we isolated eight mutants that exhibited increased resistance to serine in five distinct selection experiments. One mutant for each selection was subjected to whole genome sequencing to identify the underlying mutations. In one of the mutants (GP2324, isolated from the wild type 168 at 1 mM serine), a *ybeC* mutation was detected. Thus, we tested the remaining mutants for the presence of mutations in *ybeC*. Strikingly, four out of the eight mutants had acquired mutations in *ybeC*. These mutations resulted in the production of truncated and therefore possibly inactive YbeC proteins or in an in-frame deletion of 236 amino acids (in GP3050). The identification of multiple suppressor mutants affecting YbeC strongly supports the crucial role of YbeC in the resistance to serine.

Of the serine-resistant strains whose phenotype was not caused by a *ybeC* mutation, two strains derived from the wild type strain 168 had a duplication of the about 16 kb *yokD-thyB* chromosomal region. Interestingly, this region contains the *ilvA* gene encoding threonine dehydratase involved in the biosynthesis of isoleucine from threonine. The remaining three mutants (derived from the *serA* mutant, and the *ybeC* mutant at 10 and 17 mM serine, see Table S2 for details) had mutations affecting the repressor for the threonine biosynthetic genes, *thrR* (Rosenberg *et al*., 2016), and mutations in the regulatory regions of the *sdaAB* and *hom* promoter regions, respectively (Table S2). All these genes are involved in serine and threonine metabolism suggesting a close relation between the metabolic pathways for these similar amino acids (see below for further analyses).

### YbeC is a serine transporter

All three targeted and unbiased analyses of serine-resistance mutants identified YbeC as the main player. YbeC is similar to known amino acid transporters, is classified as a member of the amino acid-polyamine-organocation superfamily (see Table S1), and the *ybeC* mutant had the phenotypes expected for a major serine transporter. To test this idea, we made use of an *E. coli* mutant that lacks the major serine transporter SstT. This strain is less sensitive to growth inhibition by serine (Ogawa *et al*., 1997; Ogawa *et al*., 1998). We cloned the *ybeC* gene into the expression vector pWH844 and used the resulting plasmid pGP2987 to transform the *sstT* mutant JW3060 (Baba *et al*., 2006). Indeed, the expression of plasmid-borne *ybeC* in *E. coli* JW3060 restored serine toxicity (Fig. 2B). Taken together, both the genetic characterization and the functional complementation of an *E. coli* mutant lacking a serine transporter demonstrate that YbeC is indeed a transporter for serine.

The *ybeC* gene forms a monocistronic transcription unit (Nicolas *et al*., 2012). To study the activity of the *ybeC* promoter and its possible regulation by serine, a 257 bp region (222 bp upstream of the ATG translational start codon, and 35 bp of the *ybeC* coding region) was fused to a promoterless *lacZ* gene. The resulting strain, GP2965, was cultivated in minimal in the presence and absence of serine as well as in complex (LB) medium. With very similar β-galactosidase activities (144 ± 2, 132 ± 8, and 135 ± 10 units per mg of protein, respectively), this fusion was similarly expressed irrespective of the presence of serine in the medium thus indicating constitutive expression of *ybeC*.

### Serine and threonine share overlapping transporters

#### Threonine transporters contribute to serine uptake

The identification of viable *serA ybeC* mutants in the suppressor screen (Screen #2 above) suggested either that the mutant YbeC proteins retained some transport activity or that YbeC is not the only transporter for serine. To resolve this issue, we deleted the *ybeC* gene in the *serA* mutant which is auxotrophic for serine. The resulting double mutant GP2941 depends on serine uptake for viability. Analysis of growth of these mutants demonstrated that both the *serA* mutant and the *serA ybeC* double mutant were unable to grow in the absence of serine (C-glucose medium). In contrast, the strains lacking the *ybeC* gene were able to grow in minimal medium supplemented with serine (see Fig. 3A). Thus, the Δ*ybeC* mutant is still able to transport serine.

**Fig. 3.**
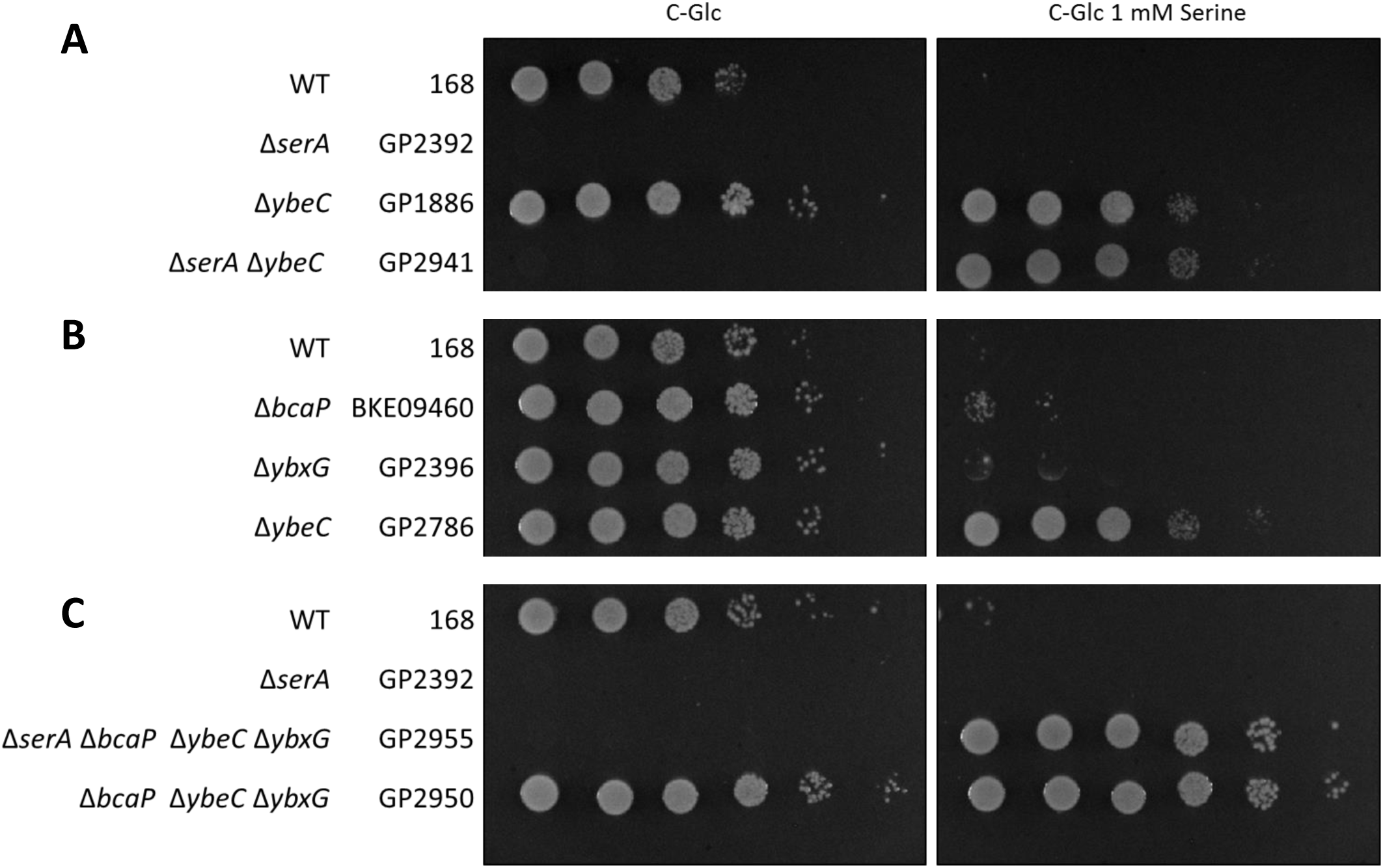
The contribution of threonine transporters to serine uptake. Cells of the indicated strains were grown in C-Glc minimal medium to an OD600 of 1.0 and serial dilutions (10-fold) were prepared. These samples were plated on C-Glc minimal plates containing no serine or 1 mM serine. The plates were incubated at 37°C for 48 h. **A.** Combination of the *ybeC* deletion with the deletion of the *serA* gene encoding phosphoglycerate dehydrogenase. The growth of the single deletion mutants of *ybeC* (GP1886) and *serA* (GP2392) was compared to the growth of the combined deletion strain of *ybeC* and *serA* (GP2941). **B.** The resistance of the threonine transporter deletion strains to serine. The *bcaP* (BKE09460) and *ybxG* deletion strains (GP2396) are compared to the wild type strain 168 and the *ybeC* deletion strain (GP1886). **C.** Combination of the *serA* deletion with the deletion strain if *bcaP, ybeC* and *ybxG*. The growth of the wild type strain 168 was compared to GP2392 (*serA*), GP2955 (*serA bcaP ybeC ybxG*) and GP2950 (*bcaP ybeC ybxG*).

Serine and threonine are chemically similar to each other and the *E. coli* SstT transporter is capable of transporting both serine and threonine (Kim *et al*., 2002). Therefore, we considered the possibility that threonine transporters might contribute to serine uptake in *B. subtilis*, and *vice versa*. Based on the analysis of growth inhibition by threonine and its analogue 4-hydroxythreonine, the BcaP and YbxG permeases have been identified as tentative threonine transporters in *B. subtilis* (see Table S1, Belitsky 2015; Commichau *et al*., 2015). To test the possible role of these permeases in serine transport, we used single, double and triple mutants lacking *ybeC, bcaP*, and *ybxG*, respectively. The resulting strains were assayed for their resistance towards serine. As shown in Fig. 3B, the single *bcaP* and *ybxG* deletions conferred only a weak resistance to growth inhibition by serine, whereas the loss of *ybeC* resulted in a substantial resistance (see also Table 2). This observation confirms that YbeC is the main transporter for serine in *B. subtilis*.

The double mutants lacking YbeC and one of the threonine transporters exhibited a substantial increase in resistance to serine indicating that both threonine permeases contribute to serine transport (see Table 2). In contrast, the *bcaP ybxG* double mutant was much more sensitive to serine than the *ybeC* mutant. This observation supports the conclusion that YbeC is the major serine permease. The analysis of double mutants lacking YbeC and either YbxG or BcaP indicates that the loss of BcaP has a higher contribution to serine resistance as compared to the loss of YbxG (Table 2, compare GP2951 and GP2949). This indicates that BcaP may be more active in serine transport than YbxG. The deletion of the three permease-encoding genes in the triple mutant GP2950 resulted in an unprecedented resistance to serine up to 100 mM (Table 2). This finding indicates that these three proteins may be responsible for the majority of serine uptake in *B. subtilis*. If these proteins were the only serine permeases, one would expect that an auxotrophic *serA* mutant lacking the three permeases would not be viable. However, this mutant (GP2955) was still able to grow on minimal medium in the presence of serine (Fig. 3C). Thus, *B. subtilis* possesses at least one additional permease that is able to transport serine.

#### Analysis of threonine transport

Our findings demonstrate that the two previously suggested threonine transporters are also active as minor serine permeases. Next, we asked whether YbeC is also capable of transporting threonine. To address this question, we made use of the observation that threonine is toxic for *B. subtilis* if added in concentrations exceeding 50 μg/ml (Lamb and Bott, 1979a; Lamb and Bott, 1979b). In our experimental setup, threonine (10 mM) inhibits growth of *B. subtilis* 168. Inactivation of the *bcaP* gene conferred a growth advantage, the *bcaP* mutant grew in the presence of threonine as well as the wild type strain in the absence of this amino acid. In contrast, the deletions of *ybxG* or *ybeC* had only minor effects (Fig. 4A). This observation is supported by the analysis of the double and triple mutants: While all mutants lacking *bcaP* showed threonine-resistant growth, this was not the case for the *ybeC ybxG* double mutant GP2952 that still expresses BcaP (Fig. 4B). These observations suggest that BcaP is the main threonine transporter in *B. subtilis*.

**Fig. 4.**
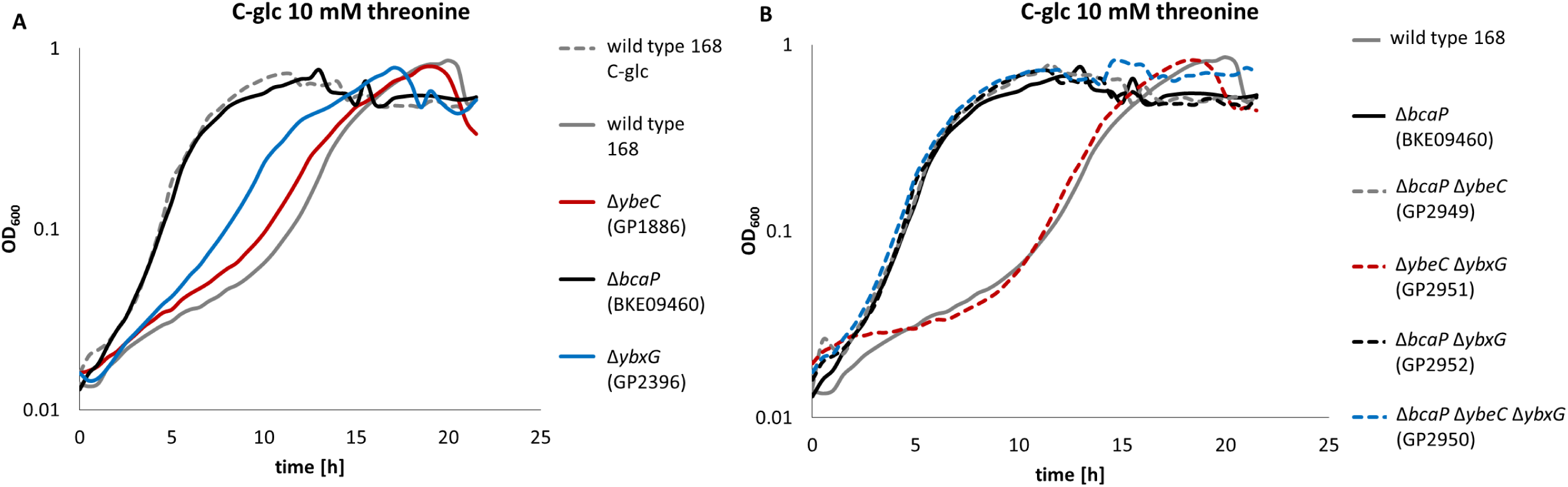
The growth inhibition by threonine. **A** The single deletion strains for *ybeC* (GP1886), *bcaP* (BKE09460) and *ybxG* (GP2396) were grown in C-glc medium with 10 mM threonine in comparison to the wild type strain 168 and the wild type strain 168 in C-glc medium without threonine. **B** The growth of the double deletion mutants *bcaP ybeC* (GP2949), *ybeC ybxG* (GP2951) and *bcaP ybxG* (GP2952) was compared to the growth of the *bcaP ybeC ybxG* deletion strain and the wild type strain 168 in C-glc medium with 10 mM threonine.

In order to test the presence of additional threonine transporters, and to get further evidence for the relative roles of BcaP, YbxG, and YbeC in threonine uptake, we deleted the *thrC* gene in the wild type 168 and in relevant transporter mutants. The *thrC* gene codes for threonine synthase which catalyzes the final step in threonine biosynthesis. As expected, the *thrC* mutant was auxotrophic for threonine (data not shown). The deletion of *bcaP* alone or in combination with *ybeC* resulted in improved growth both at 0.04 and 4 mM threonine as compared to the single *thrC* mutant (see Fig. 5). The combination of the *thrC, ybeC* and *ybxG* mutations had no effect as compared to the single *thrC* deletion supporting the idea that YbeC and YbxG play only very minor roles in threonine uptake. However, the simultaneous deletion of *bcaP* and *ybxG* in the *thrC* mutant resulted in severely reduced growth at 0.04 mM threonine (Fig. 5A). The additional deletion of *ybeC* had only a minor, if any impact. These findings suggest that BcaP and YbxG act as threonine transporters. Importantly, the *thrC* mutant lacking BcaP and YbxG (and YbeC) is still able to grow in the presence of threonine suggesting the existence of at least one additional threonine transporter (Fig. 5A and 5B).

**Fig. 5.**
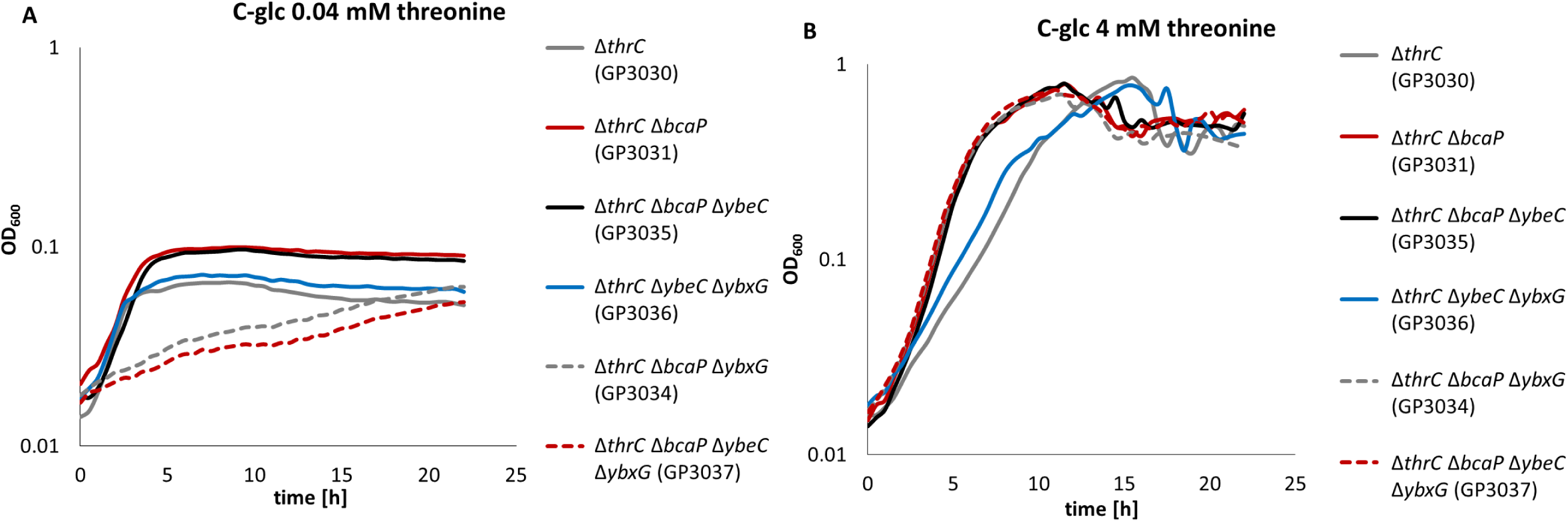
Growth of the auxotrophic strains in combination with transporter deletions in the presence of different amounts of threonine. The growth of the deletion strains GP3031 (*thrC bcaP*), GP3035 (*thrC bcaP ybeC)*, GP3036 (*thrC ybeC ybxG*), GP3034 (*thrC bcaP ybxG*) and GP3037 (*thrC bcaP ybeC ybxG*) was compared to the *thrC* deletion mutant (GP3030) in C-glc medium with 0.04 mM threonine (A) and 4 mM threonine (B).

Taken together, these results indicate that BcaP is the major threonine transporter in *B. subtilis*, whereas YbxG has a minor threonine permease activity. Our data do not support the annotation of YbeC as a threonine transporter. Moreover, BcaP and YbxG have overlapping activity for both serine and threonine (see Fig. 1).

### Putting serine toxicity in its metabolic context

As mentioned above, the addition of serine to minimal medium is toxic, whereas its addition to complex LB medium did not interfere with growth of *B. subtilis* 168 (data not shown). The inactivation of the *ybeC* gene to prevent serine uptake or the addition of several individual amino acids such as threonine overcome serine toxicity (Vandeyar and Zahler, 1986; Lachowicz *et al*., 1996). These observations suggest that intracellular serine interferes with amino acid metabolism. Mutants from the suppressor screen (Selection #2) and from the genome wide screen (Selection #3) shed light on the origins of serine toxicity.

#### The role of serine deaminase in overcoming serine toxicity

The *sdaAB-sdaAA* operon encodes the two subunits of serine deaminase, which catalyzes the degradation of serine to pyruvate and ammonia (Chen *et al*., 2012). Both the suppressor screen (Selection #2) and the whole genome screen (Selection #3) identified overexpression of *sdaAB-sdaAA* as relieving serine toxicity. In the suppressor screen, strain GP2971 had a mutation 70 bp upstream of the start of the *sdaAB* coding sequence suggesting that it might affect expression of the operon. Indeed, a promoter has been identified in the 139 bp intergenic region between the *yloV* and *sdaAB* genes (Nicolas *et al*., 2012). To test this hypothesis, we fused the 166 bp wild type and mutant regions that contain the complete *yloV*-*sdaAB* intergenic region, and thus the *sdaAB* promoter, to a promoterless *lacZ* gene, and compared the gene expression driven by these promoters. The strains carrying the *lacZ* fusions integrated into the *amyE* gene were cultivated in minimal medium, and their β-galactosidase activities were determined. For the wild type promoter, we detected 7.4 ± 2.2 units of β-galactosidase per mg of protein. This corresponds to a very weak promoter activity (Schilling *et al*., 2007). Expression of the *lacZ* gene from the mutant promoter resulted in 370 ± 48 units of β-galactosidase per mg of protein. Thus, the mutation resulted in a 50-fold increase of promoter activity. A closer inspection of the sequence around the mutation suggests that a TTGCCA sequence had been altered to the perfect −35 sequence, TTGACA. It is tempting to speculate that this perfect −35 region is responsible for the higher expression of the *sdaAB-sdaAA* operon and thus for higher intracellular levels of serine deaminase in the mutant. This conclusion is strongly supported by two mutants from the whole genome screen, affected in *yloV* (BKE15840) and *yloU* (BKE15830) that suppressed serine toxicity due to overexpression of the *sdaAB-sdaAA* operon. These strains have their antibiotic cassettes with the strong outwardly facing promoter immediately upstream of *sdaAB-sdaAA*, indicating that their overexpression suppresses serine toxicity (Table 1). The increased degradation of serine by serine deaminase is likely to be responsible for the protective action observed upon overexpression of this enzyme.

#### The role of threonine metabolism in serine toxicity

Two different loci related to threonine metabolism were identified in our screens. First, the suppressor screen identified a duplication of the 16 kb *yokD-thyB* region containing *ilvA* as relieving serine toxicity. Second, both the whole genome and suppressor screens (Selection #2, 3) identified overexpression of the *hom-thrC-thrB* operon as relieving serine toxicity.

The threonine dehydratase IlvA uses threonine in the initial step of isoleucine biosynthesis. We observed a duplication of the approximately 16 kb *yokD-thyB* genomic region encompassing *ilvA* in two suppressor strains. This observation implies that IlvA may become limiting in the presence of serine or contribute to scavenging excess serine. If IlvA is inhibited by serine, this could be compensated by overexpression of *ilvA* gene (due to genomic duplication) or by increased synthesis of ThrC with its moonlighting activity as threonine dehydratase (Skarstedt and Greer, 1973; Rosenberg *et al*., 2016) (see Fig. 1) which resumes isoleucine synthesis. However, this is unlikely since supplementation of isoleucine does not reduce serine toxicity (data not shown). Thus, it is more likely that *B. subtilis* IlvA may also have serine dehydratase activity, resulting in deamination of serine as has been shown in *Salmonella enterica* and *E. coli* (Borchert and Downs, 2018). To test whether IlvA is a major determinant for serine resistance in these suppressor strains, we overexpressed the *ilvA* gene in the wild type strain 168 using the expression vector pGP2289 (see Fig. 6). Indeed, *ilvA* overexpression provided resistance to serine. However, the level of resistance of the overexpressing strain was lower than observed for the original suppressor mutation with the genomic duplication (see Discussion).

**Fig. 6.**
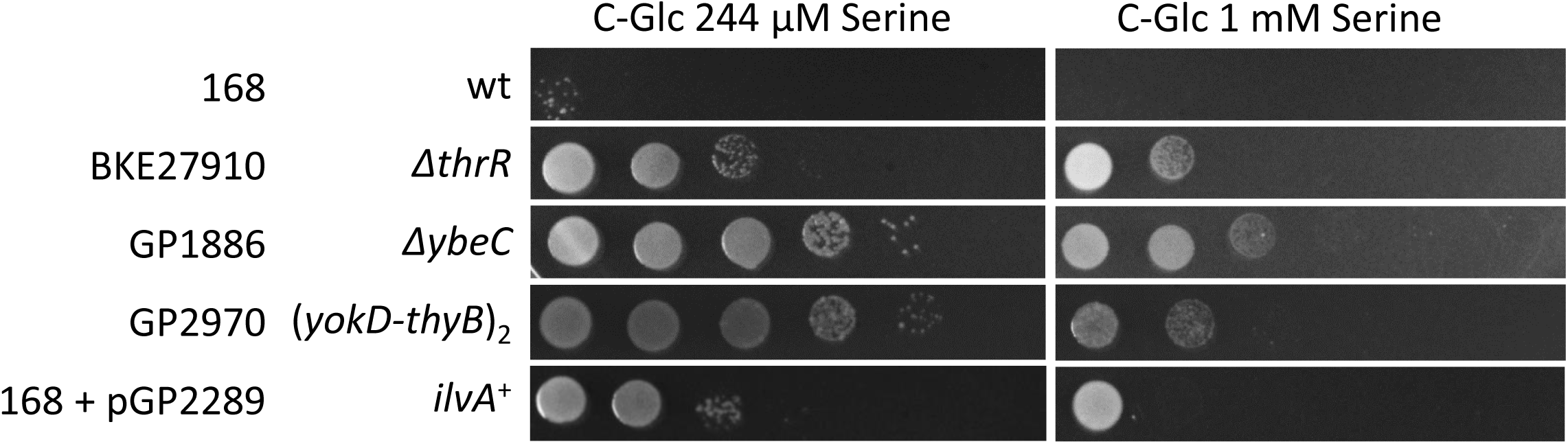
Serine resistance of the *thrR* deletion mutant and the *ilvA* overexpression strain. The growth of the wild type strain 168 and the mutant strains BKE27910 (*thrR*), GP1886 (*ybeC*), GP2970 (Suppressor with (*yokD-thyB*) duplication) and the wild type 168 with the plasmid pGP2289 (*ilvA* overexpression) was compared on C-Glc minimal medium plates (10-fold serial dilution) containing 244 µM or 1 mM serine. The plates were incubated for 48 h at 37°C.

Both the whole genome and suppressor screens identified inactivation of the ThrR repressor and overexpression of one of its target operons, *hom-thrC-thrB* as relieving serine toxicity. The suppressor screen (Selection #2) identified a mutation in *thrR* and a mutation upstream of the *hom-thrC-thrB* operon. The inspection of the mutation in the *hom* upstream region revealed that this mutation did affect the ThrR binding site (Rosenberg *et al*., 2016). Moreover, the *thrR* mutation (deletion of A91) resulted in a frame-shift and translation stop after 35 amino acids. This truncation has been observed previously in a different context. It results in an inactive ThrR protein (Rosenberg *et al*., 2016). This suggests that both the *thrR* and the *hom* promoter region mutations result in increased expression of the *hom-thrC-thrB* operon. To test this idea, we tested the activity of the wild type and mutant *hom* promoters using *hom-lacZ* fusions. Strains carrying these fusions were grown in minimal medium and their β-galactosidase activities were assayed. For the wild type promoter, we detected 275 ± 35 units of β-galactosidase per mg of protein, whereas the mutant promoter resulted in 970 ± 85 units of β-galactosidase per mg of protein. These values are similar to those determined previously for the wild type *hom* promoter and for promoter variants that carry mutations in the ThrR binding site (Rosenberg *et al*., 2016). Thus, these mutations allow an increased expression of the *hom-thrC-thrB* operon. These findings are supported by the results from the whole genome screen (Selection #3): The screen identified a strain with a *thrR* deletion and as well as overexpression of the *hom-thrC-thrB* operon originating from *yutH*, which is adjacent to the *hom-thrC-thrB* operon as relieving serine toxicity (Table 1).

Taken together, our results suggest that serine might cause defects in threonine and isoleucine biosynthesis. The defects can be overcome by reducing serine uptake, by degradation of serine, or by an adjustment of threonine and isoleucine metabolism.

## Discussion

Metabolite toxicity is one of the least understood areas in the field of microbial metabolism. However, toxic metabolites pose major problems if metabolic pathways are assembled for biotechnological applications or when approaching genome minimization (Commichau *et al*., 2015; Reuss *et al*., 2016). For *B. subtilis*, only recently significant effort has been put into the elucidation of resistance mechanisms that allow the bacterium to cope with toxic metabolic intermediates and substrates (Lambrecht *et al*., 2012; Commichau *et al*., 2015; Niehaus *et al*., 2017; Niehaus *et al*., 2018; Sachla and Helmann, 2019).

In this work, we isolated *B. subtilis* mutants that are able to grow in minimal medium supplemented with the toxic amino acid serine using three different approaches, i. e. (i) a targeted screen, (ii) an unbiased suppressor screen, and (iii) a whole genome screen. Our two laboratories initiated this project independently starting with different aims, the identification of serine transporter and understanding the origin of serine toxicity, but the information obtained from all three strategies was highly similar and complementary. The convergence of the results from the unbiased and the genome-wide screens strongly suggests that the screens were saturating and that we have elucidated the complete portfolio of possibilities that allows *B. subtilis* to cope with otherwise toxic serine concentrations.

All three different screen identified YbeC as the major serine transporter in *B. subtilis*. Moreover, this transporter works well in *E. coli* in which YbeC restores serine sensitivity of a *sstT* mutant. Three features makes the identification of amino acid transporters difficult: First, bacteria usually contain multiple transporters for one amino acid, often high and low affinity transporters that allow optimal uptake at a wide range of substrate concentrations. In *B. subtilis*, this is the case for arginine, the branched-chain amino acids, glutamine, proline, and threonine. Second, the amino acid transporters are often not highly specific, i. e. they are able to transport multiple substrates as has been shown for BcaP and GltT in *B. subtilis*. BcaP transports the branched-chain amino acids isoleucine and valine as well as threonine (Belitsky 2015; Commichau *et al*., 2015, this work), whereas GltT is involved in the uptake of aspartate, glutamate and the toxic product glyphosate (Zaprasis *et al*., 2015; Wicke *et al*., 2019). Finally, amino acid transporters are often members of families of closely related proteins, and based on sequence comparison it is often difficult to predict substrates. For example, the branched chain amino acid transporter BcaP is a paralog of the methylthioribose transporter MtrA, and the KimA protein that is member of the amino acid-polyamine-organocation (APC) superfamily (see Table S1) does actually transport potassium (Gundlach *et al*., 2017). With YbeC and serine uptake, we had to deal with all these challenges: While YbeC is the major transporter for serine, it is not the only one. Our study demonstrates that the BcaP and YbxG transporters that can transport threonine, do also contribute to serine uptake; however, their contribution is rather minor, as can be judged from the analysis of resistance of transporter mutants to serine. Even in the absence of YbeC, YbxG, and BcaP, *B. subtilis* is still able to transport serine from the medium indicating the presence of yet additional serine transporters. Moreover, all three transporters involved in serine uptake have paralogs in *B. subtilis*, which might be responsible for the residual serine uptake in the *ybeC ybxG bcaP* triple mutant that is highly resistant to serine (see Table 2). The promiscuity of amino acid transporters is important for genome minimization projects (Reuss *et al*., 2016, Reuss *et al*., 2017). For example, BcaP alone would be sufficient to transport at least four amino acids. Thus, genes encoding additional transporters for these amino acids (including *ybeC*) can be deleted as well as the corresponding biosynthetic pathways. It seems that nature has already put this reduction of amino acid acquisition to very few transporters into reality: the highly genome-reduced *Mycoplasma* species have lost the ability to produce amino acids and therefore depend completely on their uptake from the medium. Due to the fast evolution of this group of bacteria, it has so far not been possible to identify amino acid transporters based on sequence similarity. However, the independent life of the artificial genome-reduced organism *Mycoplasma mycoides* JCVI-syn3.0 (Hutchison *et al*., 2016) indicates that this minimal bacterium possesses a complete set of amino acid transporters.

The toxicity of serine can not only be mitigated by the loss of the major serine transporter, YbeC. In addition, our screens also identified other ways to cope with increased serine concentrations, *i. e*. (i) the rapid conversion of serine to other metabolites, mostly pyruvate and (ii) the overexpression of genes involved in the synthesis of threonine (the *hom-thrC-thrB* operon). The serine deaminase complex SdaAA-AB converts serine to pyruvate and ammonia, thus detoxifying excess serine as well as allowing cells to use serine as carbon and nitrogen source. It is therefore not surprising that removal of this enzyme activity was attempted to increase the yield of serine production in *E. coli* (Li *et al*., 2012). On the other hand, it was reported that serine deaminase deficiency in *E. coli* resulted in abnormal cell division even in lysogeny broth medium (Zhang and Newman, 2008). Interestingly, strains overexpressing the *sda* operon were most highly enriched from the genome-wide pool of *B. subtilis* transposon insertion mutants if the library was grown in minimal medium supplemented with toxic concentration of serine (data not shown), likely because serine uptake is not limited but rapid conversion to pyruvate and ammonia provides both carbon and nitrogen source in these strains.

There are a couple of ways in which increased expression of the *hom-thrC-thrB* operon could suppress serine toxicity. First, increased levels of threonine biosynthetic enzymes may produce more threonine. As the addition of threonine to serine-containing minimal medium can partially overcome serine toxicity, it is likely that serine addition deprives the cell of threonine, which can be overcome either by increased threonine synthesis or by external supplementation. Second, L-serine toxicity in *E. coli* works by inhibiting both the aspartate kinase and homoserine dehydrogenase activity of the fused enzyme ThrA (Costrejaen and Truffa-Bachi, 1977), and may function analogously in *B. subtilis*. Consistent with this idea, we found that supplementation of homoserine restored the growth of wild type *B. subtilis* in the presence of serine (data not shown). Biochemical analysis with purified *B. subtilis* homoserine dehydrogenase would provide clear evidence for this hypothesis. We attempted to purify *B. subtilis* homoserine dehydrogenase from *hom* overexpressing *E. coli* strain but failed to get active enzyme. It is tempting to speculate that the increased expression of *ilvA* upon the duplication the *yokD-thyB* genomic region is the major determinant for serine resistant phenotype in this suppressor mutant since we observed that overexpression of *ilvA* phenocopied it, even though only partially (Fig. 6). One explanation for the incomplete effect of IlvA overexpression is that the enzyme not only suppresses serine toxicity, but also is itself toxic to cell possibly due to the accumulation of toxic levels of 2-oxobutanoate or 2-aminoacrylate (Borchert and Downs, 2018). Strikingly, the *ilvA* gene is present in two copies in the suppressor strain whereas it is present on multiple plasmid copies and expressed from a strong constitutive promoter in the artificial overexpression system. This may be too much of a good thing!

This study provides novel insights into important aspects of serine metabolism in *B. subtilis* and into its integration into the amino acid acquisition network. This network consists not only of biosynthetic enzymes with overlapping activities but also of the transporters that are often promiscuous and transport multiple amino acids. Our work provides a starting point for further analysis of the complex and interlocking set of proteins that carry out amino acid transport in *B. subtilis*.

## Methods

### Bacterial strains and growth conditions

All *B. subtilis* strains used in this work are derived from the laboratory wild type strain 168. They are listed in Table S2. *B. subtilis* was grown in LB (Lysogeny broth) medium, SP (sporulation) medium and in C minimal medium containing glucose and ammonium as basic sources of carbon and nitrogen, respectively (Commichau *et al*., 2008). Minimal medium was supplemented with auxotrophic requirements (at 50 mg/l) and amino acids as indicated. Plates were prepared by the addition of 17 g Bacto agar/l (Difco) to the liquid medium. *E. coli* DH5α and JW3060 (Sambrook *et al*., 1989; Baba *et al*., 2006) were used for cloning and complementation experiments, respectively. JW3060 was grown in M9 minimal medium (Sambrook *et al*., 1989) with glucose (1% w/v) as the carbon source, but lacking casamino acids. Serine was added as indicated. For the determination of the tolerated serine concentrations, bacteria were grown in C glucose minimal medium to an OD600 of 1.0 and plated on C-Glc plates containing a wide range of serine concentrations (1 to 100 mM). The growth was compared after incubation of the plates at 37°C for 48 hours.

### DNA manipulation and genome sequencing

Plasmid DNA extraction from *E. coli* were performed using standard procedures (Sambrook *et al*., 1989). Restriction enzymes, T4 DNA ligase and DNA polymerases were used as recommended by the manufacturers. *Fusion* DNA polymerase (Biozym, Germany) was used for the polymerase chain reaction as recommended by the manufacturer. DNA fragments were purified using the Qiaquick PCR Purification kit (Qiagen, Germany). DNA sequences were determined using the dideoxy chain termination method (Sambrook *et al*., 1989). All plasmid inserts derived from PCR products were verified by DNA sequencing. Chromosomal DNA of *B. subtilis* was isolated as described (Commichau *et al*., 2008). To identify the mutations in the suppressor mutant strains GP2324, GP2969, GP2970, GP2971, and GP2972 (see Table S2), the genomic DNA was subjected to whole-genome sequencing (Reuß *et al*., 2019). Briefly, the reads were mapped on the reference genome of *B. subtilis* 168 (GenBank accession number: NC_000964) (Barbe *et al*., 2009). Mapping of the reads was performed using the Geneious software package (Biomatters Ltd., New Zealand) (Kearse *et al.*, 2012). Single nucleotide polymorphisms were considered as significant when the total coverage depth exceeded 25 reads with a variant frequency of ≥90%. All identified mutations were verified by PCR amplification and Sanger sequencing.

### Transformation and phenotypic analysis

Standard procedures were used to transform *E. coli* (Sambrook *et al*., 1989) and transformants were selected on LB plates containing ampicillin (100 µg/ml). *B. subtilis* was transformed with plasmid or chromosomal DNA according to the two-step protocol described previously (Kunst and Rapoport, 1995). Transformants were selected on SP plates containing chloramphenicol (Cm 5 µg/ml), kanamycin (Km 5 µg/ml), spectinomycin (Spc 150 µg/ml), or erythromycin plus lincomycin (Em 25 µg/ml and Lin 25 µg/ml).

In *B. subtilis*, amylase activity was detected after growth on plates containing nutrient broth (7.5 g/l), 17 g Bacto agar/l (Difco) and 5 g hydrolyzed starch/l (Connaught). Starch degradation was detected by sublimating iodine onto the plates.

Quantitative studies of *lacZ* expression in *B. subtilis* were performed as follows: cells were grown in LB medium or in C glucose medium supplemented with serine as indicated. Cells were harvested at OD600 of 0.6 to 0.8. β-Galactosidase specific activities were determined with cell extracts obtained by lysozyme treatment as described previously (Kunst and Rapoport, 1995). One unit of β-galactosidase is defined as the amount of enzyme which produces 1 nmol of o-nitrophenol per min at 28° C.

### Construction of deletion mutants

Deletion of amino acid transporter and biosynthetic genes was achieved by transformation with PCR products constructed using appropriate oligonucleotides to amplify DNA fragments flanking the target genes and intervening antibiotic resistance cassettes (Guerot-Fleury *et al*., 1995) as described previously (Wach, 1996).

### Whole genome growth phenotype screen

The screen was carried out as described previously (Koo *et al*., 2017) with modifications that optimized screening for serine toxicity. Plates for screening were allowed to dry for two days. The BKE (Erm^R^) library was arrayed in 384-well plates using a Biomek FX liquid handling robot (Beckman Coulter) and stored as glycerol stock. To screen the whole BKE library, cells were pinned from glycerol stocks onto rectangular LB agar plates in 384-format using a Singer Rotor robot, then four 384-format plates were combined and pinned to 1536-format. For each screen, exponentially growing cells in 1536-format were then pinned to glucose minimal agar plates (growth control) and glucose minimal plates supplemented with three different concentrations of L-serine (0.38, 0.75 and 1.5 mM). Then, plates were incubated at 37°C in a humidified incubator for about 24 to 44 hours. Plates were imaged using a Powershot G10 camera (Canon) and serine-resistant mutants were identified by their position in the plates. Each mutant was confirmed by sequencing of their barcodes.

### Plasmids

Plasmid pAC5 (Martin-Verstraete *et al*., 1992) was used to construct translational fusions of the *ybeC, sdaAB*, and *hom* control regions with the *lacZ* gene. For this purpose, the regions upstream of these genes were amplified using appropriate oligonucleotides. The PCR products were digested with *Eco*RI and *Bam*HI PCR and cloned into pAC5 linearized with the same enzymes. The resulting plasmids were pGP2287 (*ybeC*), pGP2295 (*sdaAB*), pGP2294 (*sdaAB*\*), pGP2296 (*hom*\*).

For the expression of YbeC in *E. coli*, we constructed plasmid pGP2987. For this purpose the *ybeC* gene was amplified using chromosomal DNA of *B. subtilis* as a template. The PCR product was digested with BamHI and SalI and cloned into the expression vector pWH844 (Schirmer *et al*., 1997).

For the expression of the threonine dehydratase IlvA in *B. subtilis*, plasmid pGP2289 was constructed by cloning a DNA fragment covering the *ilvA* gene between the BamHI and SalI restriction sites of the overexpression vector pBQ200 (Martin-Verstraete *et al*., 1994).

## Acknowledgements

We are grateful to Christina Herzberg for the help with some drop dilution assays and to Fabian M. Commichau for helpful discussions. This work was supported by the EU Horizons 2020 program (Rafts4Biotech, 720776 to J.S.), the Deutsche Forschungsgemeinschaft via priority program SPP1879 (to J.S.) and the NIH (R35 GM118061 to C.A.G.).

