## Supplemental Tables for "Resistance to serine in *Bacillus subtilis*: Identification of the serine transporter YbeC and of a metabolic network that links serine and threonine metabolism"

**Table S1. Known and potential amino acid transporters in *B. subtilis***

| **Transporter** | **Transporter family** | **Regulon** | **Expression^1^** | **Substrate** |
| --- | --- | --- | --- | --- |
| AapA | APC superfamily |  |  |  |
| AlsT | Alanine or glycine cation symporter family | GlnR, TnrA |  | Glutamine |
| ArtPQR | ABC transporter |  |  | Arginine |
| AzlCD |  | AzlB |  | BCAA (export) |
| BcaP | APC superfamily | CodY |  | Threonine, Isoleucine, Valine, Serine |
| BraB | BCAA transporter | CodY, ScoC |  | Isoleucine, Valine |
| BrnQ | BCAA transporter | AzlB |  | Isoleucine, Valine |
| DctP | Dicarboxylate/amino acid:cation symporter | CcpA |  | Succinate, Fumarate |
| GabP | APC superfamily | CodY, TnrA |  | Gamma-amino butyrate, Proline |
| GerAB | APC superfamily | SigF, SigG | Sporulation | Alanine |
| GerBB | APC superfamily | SigG | Sporulation | Glucose, Fructose |
| GerKB | APC superfamily | SigG | Sporulation | Asparagine, Glucose, Fructose |
| GlnHMPQ | ABC transporter | TnrA, SigE | Sporulation | Glutamine |
| GlnT | Alanine or glycine cation symporter family | GlnL | Not expressed | Glutamine |
| GltP | Dicarboxylate/amino acid:cation symporter |  |  | Glyphosate |
| GltT | Dicarboxylate/amino acid:cation symporter |  |  | Glutamate, Aspartate, Glyphosate |
| HutM | APC superfamily | CodY, HutP, CcpA |  | Histidine |
| KimA | APC superfamily | c-di-AMP |  | Potassium |
| MetNPQ | ABC transporter | CodY, S-box |  | Methionine |
| MtrA | APC superfamily |  |  | Methylthioribose |
| OpuE | Sodium-solute symporter | SigB, CcpA |  | Proline |
| PutP | Sodium-solute symporter | CodY, PutR |  | Proline |
| RocC | APC superfamily | AhrC, CodY, RocR |  | Arginine |
| RocE | APC superfamily | AhrC, CodY, RocR |  | Arginine |
| SteT | APC superfamily |  |  | Serine/ threonine exchange |
| TcyABC | ABC transporter |  |  | Cystine |
| TcyJKLMN | ABC transporter | CymR, AscR |  | Cystine |
| TcyP | Dicarboxylate/amino acid:cation symporter | CymR |  | Cystine |
| TrpP | ECF transporter | MtrB |  | Tryptophan |
| YbeC | APC superfamily |  |  | Serine |
| YbgF | APC superfamily |  |  |  |
| YbxG | APC superfamily |  |  | Threonine, Serine |
| YcgH | APC superfamily |  | Sporulation |  |
| YdgF | APC superfamily |  |  |  |
| YecA | APC superfamily |  |  |  |
| YfkT | APC superfamily | SigG | Sporulation | Germinants |
| YflA | Alanine or glycine cation symporter family | SigB | Stress |  |
| YhjB | Sodium-solute symporter | CodY |  |  |
| YndE | APC superfamily | SigG | Sporulation | Germinants |
| YodF | Sodium-solute symporter | TnrA |  |  |
| YrbD | Alanine or glycine cation symporter family | TnrA | Sporulation | Alanine |
| YtnA | APC superfamily |  |  | Alanine |
| YvbW | APC superfamily | T-box |  | Leucine |
| YveA | APC superfamily | SigG, DegU | Sporulation |  |
| YvsH | APC superfamily | L-box |  | Lysine |
| YwcA | Sodium-solute symporter | SigE | Sporulation | Acetate |
| YxeMNO | ABC transporter | CymR |  | S-(2-succino)cysteine |

**^1^** Expression is indicated for potential transporters that are not expressed during vegetative growth suggesting that they are not playing a role in amino acid transport under standard growth conditions.

**Table S2. *Bacillus subtilis* strains used in this study.**

| Strain ^1^ | Genotype | Source ^2^ |
| --- | --- | --- |
| 168 | *trpC2* | Laboratory collection |
| BKE02120 | *trpC2 ΔybeC::erm* | Koo *et al*., 2017 |
| BKE02130 | *trpC2 ΔglpQ::erm* | Koo *et al*., 2017 |
| BKE09460 | *trpC2 ΔbcaP::erm* | Koo *et al*., 2017 |
| BKE15830 | *trpC2 ΔyloU::erm* | Koo *et al*., 2017 |
| BKE15840 | *trpC2 ΔyloV::erm* | Koo *et al*., 2017 |
| BKE27910 | *trpC2 ΔthrR::erm* | Koo *et al*., 2017 |
| BKE32270 | *trpC2 ΔyutH::erm* | Koo *et al*., 2017 |
| BP558 | *trpC2 amyE*::(P*_hom_-lacZ cat*) | Rosenberg *et al*., 2016 |
| GP1885* | *trpC2 ΔytnA::spec* | LFH → 168 |
| GP1886* | *trpC2 ΔybeC::cat* | LFH → 168 |
| GP1887* | *trpC2 ΔyodF::cat* | LFH → 168 |
| GP1888* | *trpC2 ΔalsT::tet* | LFH → 168 |
| GP2324 | *trpC2 ybeC* (Δbp 340, stop aa 125) (*yokD-thyB*)_2_ | Suppressor from 168 |
| GP2325 | *trpC2 ybeC* (Δbp 974, stop aa 350) | Suppressor from 168 |
| GP2377* | *trpC2 ΔaapA::tet* | LFH → 168 |
| GP2378* | *trpC2 ΔsteT::cat* | LFH → 168 |
| GP2379* | *trpC2 ΔmtrA::kan* | LFH → 168 |
| GP2392 | *trpC2 ΔserA::zeo* | LFH → 168 |
| GP2395* | *trpC2 ΔyhjB::tet* | LFH → 168 |
| GP2396 | *trpC2 ΔybxG::cat* | LFH → 168 |
| GP2786 | *trpC2 ΔybeC::kan* | LFH → 168 |
| GP2799* | *trpC2 ΔgltP::kan* | LFH → 168 |
| GP2930* | *trpC2 ΔydgF::cat* | LFH → 168 |
| GP2941 | *trpC2 ∆serA::zeo ∆ybeC::cat* | GP1886 → GP2392 |
| GP2949 | *trpC2 ∆bcaP::erm ∆ybeC::kan* | BKE09460 → GP2786 |
| GP2950 | *trpC2 ∆bcaP::erm ∆ybeC::kan ∆ybxG::cat* | GP2396 → GP2949 |
| GP2951 | *trpC2 ∆ybeC::kan ∆ybxG::cat* | GP2396 → GP2786 |
| GP2952 | *trpC2 ∆bcaP::erm ∆ybxG::cat* | GP2396 → BKE09460 |
| GP2955 | *trpC2 ∆serA::zeo ∆bcaP::erm ∆ybeC::kan ∆ybxG::cat* | GP2392 → GP2950 |
| GP2965 | *trpC2 amyE*::(P*_ybeC_-lacZ cat*) | pGP2287 → 168 |
| GP2966 | *trpC2 amyE*::(P*_sdaAB_* C70A-lacZ cat*) | pGP2294 → 168 |
| GP2967 | *trpC2 amyE*::(P*_sdaAB_-lacZ cat*) | pGP2295 → 168 |
| GP2968 | *trpC2 amyE::*(P*_hom_* G56T-lacZ cat*) | pGP2296 → 168 |
| GP2969 | *trpC2 ∆serA::zeo; thrR* (Δbp 90, stop aa 36) | Suppressor from GP2392 |
| GP2970 | *trpC2* duplication (*yokD-thyB*)_2_ | Suppressor from 168 |
| GP2971 | *trpC2 ∆ybeC::kan* Promoter *sdaAB** [bp -70 C→A] | Suppressor from GP2786 |
| GP2972 | *trpC2 ∆ybeC::kan* Promoter *hom** [bp -56 G→T] | Suppressor from GP2786 |
| GP3030 | *trpC2 ∆thrC::spec* | LFH → 168 |
| GP3031 | *trpC2 ΔthrC::spec ΔbcaP::erm* | GP3030 → BKE09460 |
| GP3034 | *trpC2 ΔthrC::spec ΔbcaP::erm ΔybxG::cat* | GP3030 →GP2952 |
| GP3035 | *trpC2 ΔthrC::spec ΔbcaP::erm ΔybeC::kan* | GP3030 →GP2949 |
| GP3036 | *trpC2 ΔthrC::spec ΔybeC::kan ΔybxG::cat* | GP3030 →GP2951 |
| GP3037 | *trpC2 ΔthrC::spec ΔbcaP::erm ΔybeC::kan ΔybxG::cat* | GP3030 →GP2950 |
| GP3039* | *trpC2 ΔyecA::cat* | LFH → 168 |
| GP3040* | *trpC2 ΔybgF::cat* | LFH → 168 |
| GP3049 | *trpC2 ∆serA::zeo ybeC* (bp 1564 G→T, stop aa 522) | Suppressor from GP2392 |
| GP3050 | *trpC2 ∆serA::zeo ybeC* (Δbp 307-1014) | Suppressor from GP2392 |

**^1^** The strains labelled with an asterisk were used for the targeted transporter screen (screen #1).

**^2^** Arrows indicate construction by transformation. LFH, construction by long-flanking homology PCR using upstream and downstream fragments flanking a resistance gene.
